# *Starship* transposons mobilise via a circular intermediate with potential biotechnological applications

**DOI:** 10.64898/2026.09.21.753150

**Authors:** Allira Baericke, Vanta Jameson, Andrew S Urquhart

## Abstract

Mobile genetic elements drive genome evolution and horizontal gene transfer, yet the mechanisms underlying the mobility of many eukaryotic elements remain poorly understood. Here we show that *Starships*, a recently discovered class of giant transposable elements in fungi, mobilise via an extrachromosomal circular intermediate. Using deep sequencing, we detect low-abundance reads spanning the hypothesised junction consistent with circularisation. We demonstrate that a synthetic circular intermediate, or “*starmid*”, is sufficient for recombinase-dependent integration into the host genome and that circularisation enhances expression of the *starship*-encoded tyrosine recombinase through juxtaposition of 3′ and 5′ terminal sequences. This topology-dependent regulation parallels bacterial integrative elements, in which excision activates recombination machinery. Exploiting this circular intermediate, we explore *starmids* as a platform for transient DNA delivery, enabling CRISPR–Cas9-mediated genome editing in *Paecilomyces variotii* with subsequent curing of the construct. Together, our results shed light on the biology of *starship* mobility and suggest promising biotechnology applications for these elements.

## Introduction

Mobile genetic elements (MGEs) are pervasive drivers of genome evolution with profound medical and economic consequences ^1^. For example, mobile plasmids in bacteria underpin the rapid spread of antibiotic resistance, posing a growing global health threat ^2^. In humans, MGEs have left an equally deep imprint: inactive transposable elements constitute ∼50% of the genome, and retroviruses like HIV cause devastating pandemics by integrating into our DNA ^3,4^. On the other hand, understanding MGEs and corresponding host defences has been central to the development of molecular biology as a discipline. DNA cloning using plasmid vectors was adapted from natural mobile plasmids ^5,6^, and reverse transcriptase was derived from a retrovirus ^7,8^. T4 ligase ^9^ and cre recombinase ^10^ were both isolated from bacteriophages. Restriction enzymes ^11^ and CRISPR systems ^12^ were both discovered as prokaryotic defences against viruses. All of these discoveries have become crucial to molecular biology. This outsized influence of MGEs is reflected in at least nine Nobel Prizes awarded for related discoveries. Yet much remains to be understood about these diverse and fascinating elements.

In eukaryotes, our understanding of MGEs lags behind that in bacteria, despite the first MGEs identified in any organism being described in wheat by Barbara McClintock over 75 years ago ^13^. This holds true for the fungal kingdom, where the evolutionary and economic impacts of MGEs have only recently come into focus. This has accelerated with the discovery of *Starship* transposons ^14–16^. *Starships* are massive transposable elements (potentially exceeding 1 Mb) which uniquely among known eukaryotic mobile elements are capable of carrying significant amounts of genetic cargo both within and between genomes, including horizontally between species separated by at least 100 million years of evolution ^17^. Cargo already known to be carried on *Starships* includes genes involved in pathogenicity ^18–21^, stress tolerance ^14,22^ and domestication of fungi ^23^. There are still many gaps in our understanding of *Starships*. In particular, little is known about how *Starships* mobilise or how they might, like other MGEs, be harnessed for molecular engineering.

The mobility of *Starships* relies on the activity of a tyrosine DNA recombinase (YR) enzyme ^15^. In this sense they appear similar to the crypton transposons discovered more than two decades ago first in fungi, and then subsequently in diverse eukaryotes ^24,25^. In understanding the mobility of YR-encoding eukaryotic elements, researchers have looked towards bacterial recombinase-encoding MGEs. Significant structural similarity has been reported between the “captain” recombinases (the enzymes responsible for *Starship* transposition) and Cre recombinase ^15^. Cre recombinase’s natural function is as a site-specific resolvase in bacteriophage P1, converting circular plasmid multimers into monomers via recombination at *loxP* sites ^10^. This recombinase has found widespread applications in molecular biology, for example in the creation of tissue-specific knockouts in mice and can catalyse the formation of circular DNA molecules from linear substrates in engineered systems ^26^. While distinctly different, the most informative evolutionary analogues of *Starships* are likely to be the integrative and conjugative elements (ICEs) of bacteria ^14,27^. These elements are mobilised by a diversity of mechanisms, but the most relevant to this study are those employing tyrosine recombinases. In such systems, site-specific recombination between the flanking attachment sites of an integrated element generates a circular extrachromosomal intermediate ^27^. As such it has previously been hypothesised that *Starships* (along with *Cryptons*) might also mobilise via a circular intermediate ^15,28^. However, thus far initial attempts to find evidence of such molecules via PCR have not been successful ^18^. Here, we both test this hypothesis and explore how such intermediates might be harnessed for genetic engineering.

## Results

### Deep *Illumina* coverage reveals presence of a circular intermediate

We, along with others, have hypothesised that *Starships* move via a circular intermediate formed by recombination between the regions of microhomology (or DRs) which flank them (Figure 1A). The most extensively characterised *Starship* to date is *Hephaestus* found within *Paecilomyces variotii* ^14,15,17^. We have previously experimentally demonstrated the movement of *Hephaestus* within and between *P. variotii* strains. As such the *P. variotii*–*Hephaestus* system was an ideal model in which to search for circular *Starship* intermediates. *Starship* excision is expected to be rare, and any extrachromosomal intermediate very transient. We therefore reasoned that detection of a circular intermediate would require deep sequencing coverage. To test this, we analysed a 258.9 Gbp Illumina dataset (∼8,000× coverage) compiled from *Hephaestus*-positive strains of *P. variotii*, *P. paravariotii*, and *Aspergillus fumigatus* ^15,17,29,30^. These include wildtype *P. variotii* and *P. paravariotii* strains carrying *Hephaestus, P. variotii* strains modified to carry *Hephaestus* within an antibiotic resistance cassette, or recipient strains from experimental transfer experiments.

**Figure 1).**
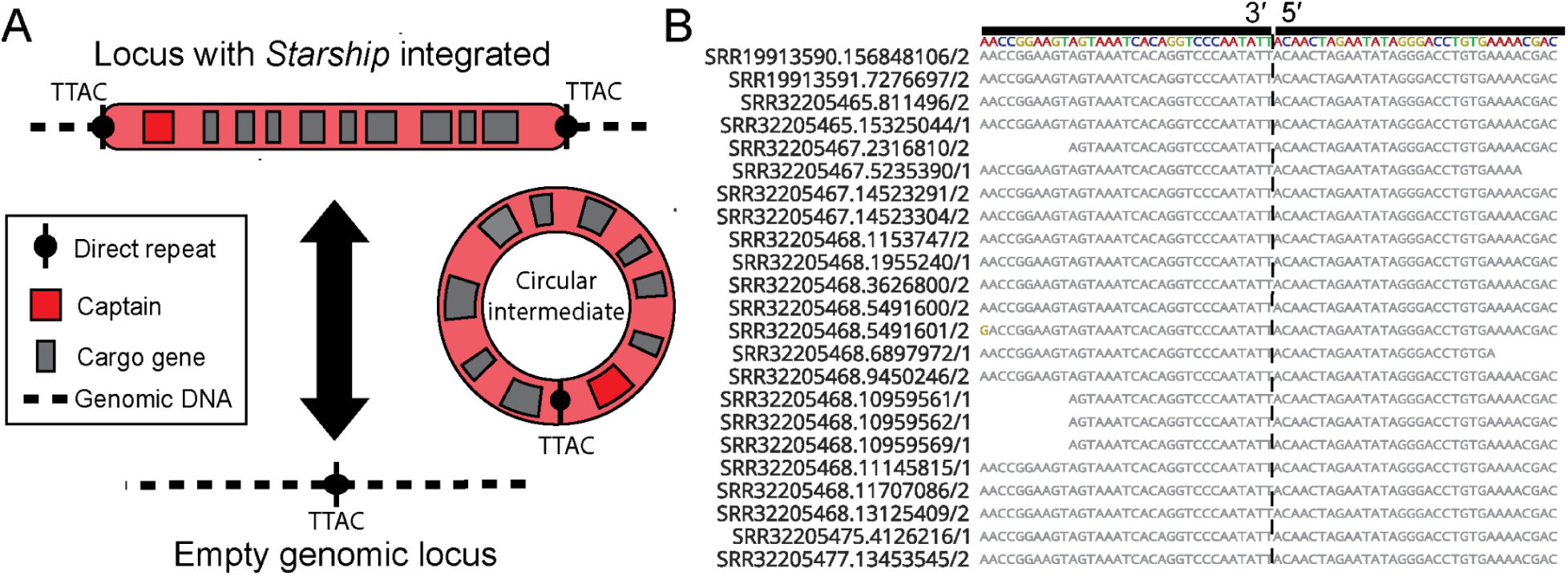
*Hephaestus* forms a circular intermediate detectable within Illumina sequencing. **A)** Diagram demonstrating the theoretical mechanism of *Starship* movement via a circular intermediate. Adapted from Urquhart et al. ^31^ **B)** Illumina sequencing reads map across the junction point on the circular intermediate form of *Hephaestus* which is formed by joining the 5′ and 3′ ends of the *Starship* (a full alignment with all reads is shown in Figure S1)

We searched for reads spanning the predicted recombination junction, defined by a 40-mer extending 20 bp either side of the expected TT//AC recombination site. In total, 36 such reads were detected across 13 separate SRA accessions (Figure 1B). In contrast, 6,959 reads contained the 3′ terminal 40-mer of *Hephaestus*. This shows that the circular intermediate exists at a low but detectable frequency, confirming that this molecule is naturally created within *Hephaestus+* strains.

### A synthetic circular intermediate is capable of self-integration into the fungal genome

Having established the creation of a circular intermediate within fungal cells, we next sought to demonstrate that this molecule is capable of reintegration into the fungal genome mediated by the captain tyrosine recombinase HhpA which we have previously demonstrated as being required for transposition ^15^. To do this we created a “*starmid*”, i.e. a plasmid vector containing the proposed circular intermediate region (Fig 2A) and transformed this directly into protoplasts of a *Hephaestus*-negative *P. variotii* strain. As a control we made two otherwise identical constructs, one containing a frameshift mutation within the *hhpA* gene. Transformation using the construct with an intact *hhpA* gene gave a consistently higher rate of transformation using equivalent quantities of plasmid DNA (mean 32.3 versus 0.83 colonies; *P* = 0.0022, two-sided Mann–Whitney U test; Figure 2B). That this integration was mediated by the *Starship’s* captain recombinase was confirmed through Illumina sequencing of a pool of transformants, which demonstrated that integration occurred at HhpA’s expected target site (Figure 2C).

**Figure 2).**
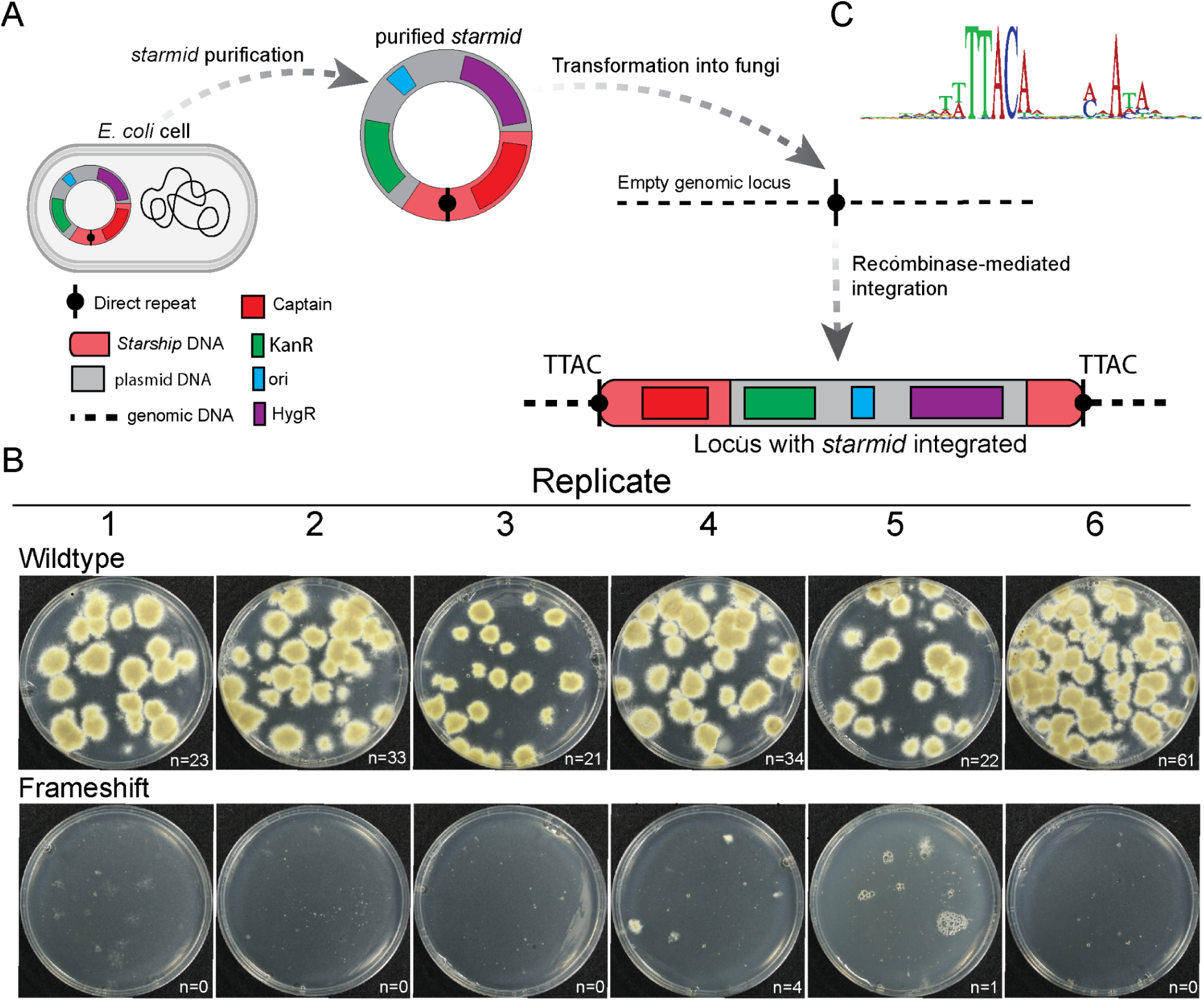
A synthetic circular intermediate is capable of self-integration into the fungal genome. **A)** A *starmid* fuses the components of a classic plasmid allowing cloning in *E. coli* (AmpR, origin of replication (ori)), a marker allowing selection in fungi (in this case HygR - a hygromycin resistance cassette), and a segment of the transient circular *Starship* intermediate including the recombination point and the *hhpA* recombinase gene. **B)** Transformation results from *starmids* with either a functional *hhpA* sequence or a *hhpA* sequence that has been inactivated with a frameshift mutation. **C)** Integration sites of the *starmid* construct within the fungal genome based on 99 integration sites.

### Circularisation enhances captain recombinase expression

Given that the proposed circular intermediate was capable of expressing the recombinase required for its own integration, we hypothesised that the circularised form of the *Starship* would promote greater captain expression than the linear form. To test this hypothesis, we cloned 10 constructs, the first of which expressed GFP under the control of the 451 bp of DNA present in the linear form. The remaining constructs added varying lengths of additional DNA corresponding to the longer circular form of the promoter (99 bp to 1,489 bp of additional DNA). Three transformants of each of these constructs (randomly integrated into the genome) were analysed for GFP expression by flow cytometry. The constructs which contained 699 bp, 899 bp, 1,099 bp, and 1,489 bp of additional sequence were shown to have a statistically significant increase in GFP expression compared to the shorter promoter sequences (Figure 3).

**Figure 3).**
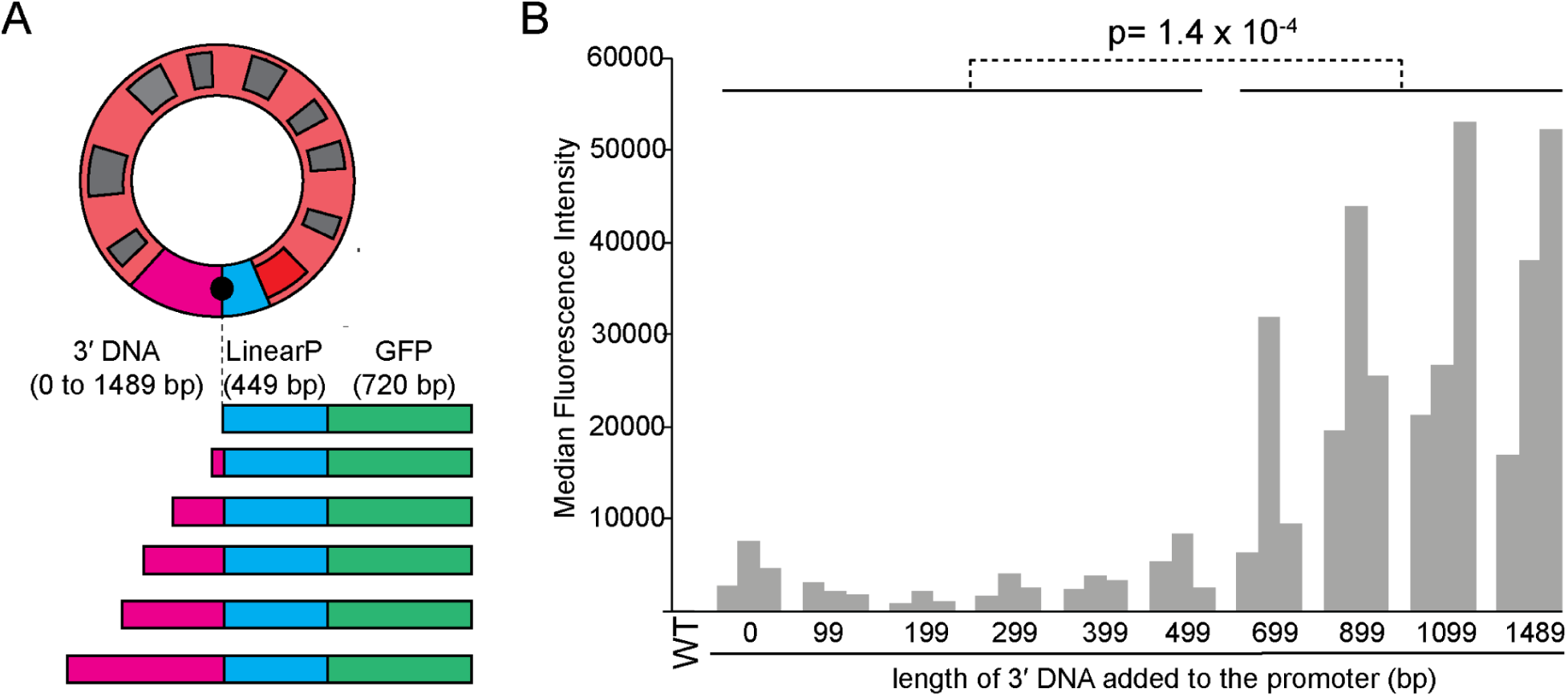
The 3′ sequence of the *Starship* forms part of the Captain’s promoter in the circular form. **A)** A series of constructs were produced in which GFP was expressed under the control of the *Starship* DNA upstream of the Captain in the linear form (“LinearP”). **B)** GFP fluorescence in conidiospores from isolates transformed with each construct measured by flow cytometry. Three independent transformants were assayed for each construct. *P* = 1.4 × 10⁻⁴, two-sided unpaired t-test.

### *Starmids* enable transient CRISPR–Cas9 delivery in fungi

A functional property of *Starship* elements is that they are semi-stable within fungal genomes, with the loss of *Hephaestus* occurring at a frequency on the order of 1 in 10^7^ *P. variotii* conidia ^15^. As such we reasoned that *Starships* might be a convenient means of temporary DNA delivery into fungal strains. As a proof of principle, we decided to test transient transformation of a gRNA/Cas9 construct on a *starmid* into *P. variotii*. The construct used had the following components within a *starmid*: a hygromycin selectable marker, CRISPR–Cas9 machinery and a thymidine kinase (TK) cassette (Figure 4A). The gene that we chose to target for CRISPR mutation was the *pvpP* gene required for pigment biosynthesis given the white phenotype that results when this gene is deleted ^32^. The TK cassette converts fluorodeoxyuridine (FdU) into a toxic product and thus can be used as a negative selectable marker in a variety of systems including fungi ^33^.

**Figure 4).**
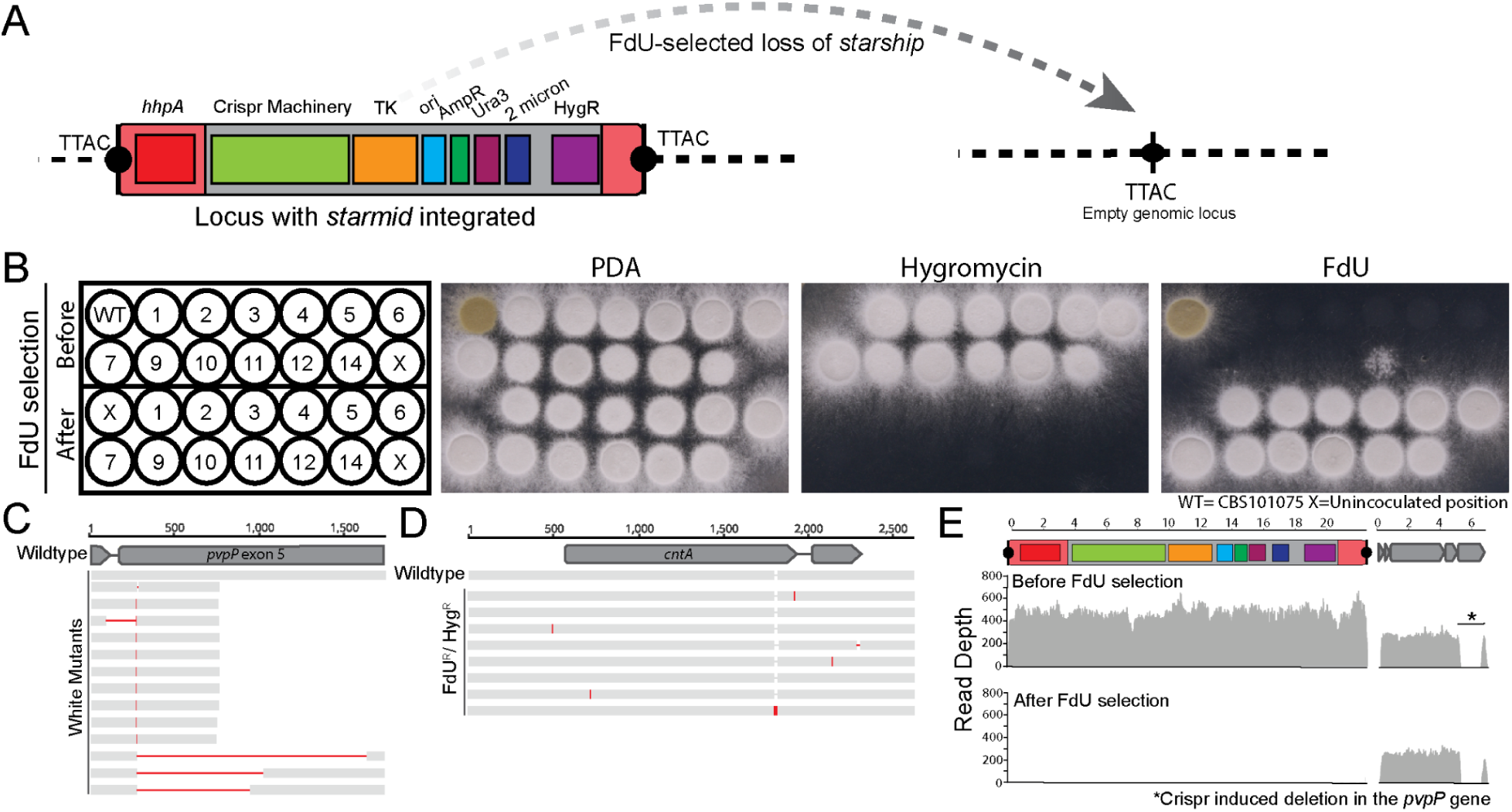
Use of a *starmid* for the temporary delivery of CRISPR–Cas9 machinery. **A)** Map of the *starmid* used for CRISPR mutation showing the additional CRISPR and thymidine kinase components **B)** Mutant phenotypes on potato dextrose agar (PDA) and on PDA with hygromycin or FdU added. White colonies indicate CRISPR–mediated disruption of the *pvpP* gene. **C)** Amplicon sequencing revealed a series of mutations within the *pvpP* gene at the target locus. **D)** Sequencing of the *cntA* gene from isolates which had gained FdU resistance without loss of hygromycin resistance (“Fdu^R^/Hyg^R^”) **E)** Illumina sequencing coverage before and after FdU selection for the loss of the TK gene.

Transformation of the CRISPR–Cas9 *starmid* into *P. variotii* resulted in the emergence of white colonies or sectors. Single spore isolation then resulted in a set of uniformly white mutants (Figure 4B). The sequencing of 14 independent white mutants showed mutations within the *pvpP* gene at the targeted site ranging from single base pair insertions to a 1.3 kb deletion (Figure 4C).

To determine whether the loss of the *starmid* could be selected for using the TK marker, we replated onto FdU and picked spontaneously resistant colonies. We then tested these on hygromycin to confirm the loss of the *starmid* construct. Unexpectedly, a number of FdU-resistant colonies retained hygromycin resistance. To understand the genetic basis for this, we performed Illumina sequencing on one such colony, and through manual inspection of potential SNPs identified a promising candidate SNP in the gene encoding NCBI XP_028486326. This gene is predicted to encode the homolog of *cntA* in *Aspergillus nidulans* which is a nucleoside transporter whose mutation confers 5-fluorouridine resistance ^34^. Amplicon sequencing of a PCR product containing the *cntA* gene identified mutations in 7 of 8 isolates tested, confirming that mutations within this gene are a dominant mode of escape from FdU selection (Figure 4D). The identification of this gene may provide avenues to improve the efficiency of FdU selection in fungi, or future studies might choose an alternative selectable marker. Nonetheless, we were able to obtain a hygromycin-sensitive, FdU-resistant spontaneous mutant from 12 of the 14 white mutants (Figure 4B).

To confirm that the hygromycin-sensitive, FdU-resistant isolates had lost the entire *starmid* construct, we Illumina sequenced an isolate before and after FdU selection. In the strain sequenced before FdU selection we saw two copies of the *starmid* construct integrated based on read depth. Following selection on FdU, both copies of this *starmid* were lost as confirmed by Illumina sequencing (Figure 4E). This demonstrates that *starmids* can effectively be used for the temporary delivery of CRISPR–Cas9 machinery.

### *Starship* excision events are not independent

Unexpectedly, we observed complete loss of multiple integrated *starmid* copies following selection. Given the estimated excision frequency (∼1 in 10⁷ conidia), independent loss of three copies would be vanishingly unlikely (∼1 in 10²¹). This suggested to us that excision events may be coordinated or coupled. To test this hypothesis, we crossed a previously-generated strain containing a miniature version of *Hephaestus* within a hygromycin resistance cassette to a wildtype strain containing the full-length *Hephaestus* and selected a progeny strain that contained both the mini-*Hephaestus* and the full-length *Hephaestus* ^15^. We then selected for the transposition of the mini-*Hephaestus* using hygromycin selection and assayed the transposition of the full-length copy of *Hephaestus* via PCR. Of 184 such strains assayed, the full-length copy of *Hephaestus* was excised in 8 (i.e. 4.3%). This is orders of magnitude higher than the expected rate of *Hephaestus* excision.

## Discussion

Our results provide two key discoveries in the emerging field of *Starship* biology: (1) that *Starship* elements mobilise via a circular intermediate, and (2) that these intermediates modified into *starmids* have research and biotechnological applications. The identification of a circular intermediate has implications for understanding how these elements move horizontally between species of fungi. Circular DNA molecules are expected to be more stable than linear fragments because they lack free ends that can be exposed to exonuclease degradation. Furthermore, we have shown here that when introduced into a new host the circular intermediate is able to produce the captain recombinase enzyme and integrate itself into the genome.

Mechanistically, *Starship* mobilisation shows striking parallels to bacterial mobile genetic elements. Our data indicate that circularisation not only enables recombination but also alters expression of the captain recombinase, with the 3′ terminal sequence contributing to activation of the 5′-encoded captain recombinase. Circularisation-dependent activation of recombinase expression has been described in bacterial integrative elements. In ICEclc^B^^13^, excision places a strong promoter upstream of the integrase gene, increasing *int* expression relative to the integrated state, in which it is driven by a weak, regulated promoter ^35^. A frequent feature of known *Starships* is the presence of an inward-facing tyrosine recombinase gene, termed the captain, at the 5′ end. Given our results, we suggest that differential expression of the captain between linear and circular configurations is one advantage of this architecture. The increased expression of the captain in the circular form of the *Starship* is one potential explanation for the higher-than-expected frequency of double-excision events in strains carrying two copies of *Hephaestus.* However, it is plausible that an unknown signal/event independently triggers both *Starship* copies to jump. Alternatively, an unknown signal or cellular state may independently trigger excision of both *Starship* copies. This would be consistent with the previous observation that the *Starships Hephaestus* and *Pegasus* can be co-transferred between strains ^17^.

Beyond their biological significance, our findings highlight the potential of *Starships* as tools for molecular engineering. The *Hephaestus Starship* targets a relatively simple consensus sequence that will be present in any potential host genome, raising the possibility of integration without prior genome modification. This distinguishes *Starships* from Cre recombinase-based systems used to introduce large regions of DNA into fungi which rely on the introduction of a “landing pad” ^36^. The largest *starmid* in this study already introduced 23 kbp of DNA and the upper limit for how much DNA a single *starmid* could be used to introduce remains to be explored. *Starships* are a remarkably diverse class of elements and target a varied range of genomic integration sites. As such, *starmids* could be readily designed to target a number of different integration sites including ones with greater target site specificity ^37^.

A particular strength of *starmids* is that they can be used for scarless transient delivery of DNA constructs. Stable plasmids are only available for use in a relatively restricted range of filamentous ascomycetes, notably the AMA1-based plasmids used extensively in *Aspergillus* ^38^. In comparison to the AMA1 plasmids the *starmids* are more stable requiring counter-selection to cure strains, as opposed to AMA1 plasmids which are rapidly lost without selection ^38^. We demonstrated the successful CRISPR mutation of *pvpP* in *P. variotii* using our *starmid* system. CRISPR mutations have previously been made in *P. variotii* using both plasmid-encoded CRISPR/Cas9-gRNA and *in vitro* assembled CRISPR/Cas9-gRNA complexes ^39,40^. *Starmids* offer advantages over currently available tools in *Paecilomyces:* an AMA1 plasmid-based system had the disadvantage of frequent integration of parts of the plasmid construct into the host genome and editing with *in vitro*-assembled CRISPR/Cas9-gRNA complexes has yet to be demonstrated without applying selection. Editing with CRISPR/Cas9-gRNA can be sufficiently efficient to negate the need for selection as has been demonstrated in *Aspergillus fumigatus* but this might require extensive optimisation for each species ^41^. For those species where protoplasting protocols are not available or are challenging, *Agrobacterium*-mediated transformation could be used to introduce the CRISPR–Cas9 components within a miniature linear *Starship* which could then be removed via FdU selection, although with the disadvantage that a scar (containing the right and left T-DNA borders) would remain at the site of T-DNA integration.

In the longer term, the full biotechnological impact of *starmids* will be unlocked if *starmids* can be first transformed into an experimentally amenable species such as *P. variotii* and then subsequently moved via horizontal *Starship* transfer into less experimentally amenable hosts. This would open up a vast number of fungal species to molecular engineering for which transformation methods have not yet been developed.

## Methods

### Detection of the circular intermediate within Illumina data

To identify reads corresponding to the hypothetical circular intermediate, 39 SRA accessions (Table S1) derived from strains containing *Hephaestus* were initially mapped against the hypothetical circular intermediate using Bowtie2 ^42^. The mapped reads were searched for reads containing the 40 bp sequence AAATCACAGGTCCCAATA<u>TTAC</u>AACTAGAATATAGGGACC which overlaps the expected recombination point within the TTAC flanking microhomology. These reads were then remapped to the junction sequence in Geneious Prime to produce the alignment in Figure 1B. Importantly, all sequencing data examined were generated before the artificial circular intermediate had been cloned, precluding the possibility of cross-contamination.

### Strains

*Paecilomyces variotii* wildtype strains CBS 144490 (*Hephaestus*+) and CBS 101075 (*Hephaestus*-) were used. *Agrobacterium* strain EHA105 was used for *Agrobacterium*-mediated transformation. *S. cerevisiae* BY4742 and *E. coli* DH5α were used for cloning.

### Construction of *starmid* vectors

A *starmid* was constructed which contained a hygromycin resistance cassette and a 5.7 kb fragment of the *Hephaestus* circular intermediate comprising 1,887 bp of 3′ sequence and 3,832 bp of 5′ sequence (including the captain recombinase gene *hhpA*). The hygromycin resistance cassette was amplified from plasmid PMAI6 ^43^ with primers AD31 and AD32 (Primer sequences are given in Table S2). The 3′ and 5′ ends of *Hephaestus* were amplified with primers AD33+AD34 and AD35+AD36 from both the wildtype and *hhpA*-frameshift versions of a previously published mini-*Hephaestus* construct ^15^. The three PCR fragments were combined into the plasmid pYES2 linearised by HindIII and EcoRI using homologous recombination in *S. cerevisiae*.

A *starmid* to mutate the *pvpP* gene required for conidial pigment biosynthesis was constructed containing the following additional components: the gRNA - Cas9 machinery and a thymidine kinase gene which is lethal when the medium is supplemented with the nucleoside analogue FdU ^33^. The design of the components to express the required gRNA and Cas9 proteins was based on plasmid pFC334 ^44^. The 20 bp target sequence in pFC334 was replaced with a target sequence previously used to mutate the *pvpP* gene of *P. variotii* (GGCTTCTCGACATTGATCGG) ^39^. To accomplish this, three fragments of pFC334 were amplified with primers AD190+AD191, AD192+AD193 and AD194+AD195. In addition, the gRNA target sequence with required homology for yeast-based cloning was ordered as a synthetic DNA from IDT. The thymidine kinase coding sequence was amplified using primers AD116 and AD117 from plasmid pPZPtk8.10 ^33^ and cloned into plasmid PLAU2 which contains an actin promoter derived from *Leptosphearia maculans* which we have used previously in *P. variotii* ^43^. The entire promoter-TK-terminator cassette was then amplified from this intermediate construct with primers AD196 and AD197. These five fragments were cloned into the NotI site of the basic *starmid* vector via homologous recombination in *S. cerevisiae*.

### Transformation of *P. variotii* with *starmids* and determination of the resulting genomic integration sites

One microgram of purified *starmid* was transformed into *P. variotii* protoplasts following the protocol of Arentshorst et al except that the 200 mg of lysing enzyme was replaced with 500 mg of VinoTaste Pro ^45^. Integration sites were determined via a pooled sequencing approach. Transformants from 6 transformation plates were moved directly (via scraping across the entire plates) into liquid media and grown overnight at 37°C in PDB with shaking before DNA was extracted using a CTAB-based protocol as described previously ^46^. Genomic DNA was sequenced at Victorian Clinical Genetics Services, Australia.

### GFP reporter constructs

A construct containing GFP expressed under the linear promoter was cloned by amplification of three PCR products. The hygromycin resistance cassette was amplified with primer pair AD90+AD91 from plasmid PMAI6, AD92+AD93 amplified the linear promoter of *Hephaestus* and AD94+AD95 amplified the GFP coding region and terminator of plasmid PLAU17. The three amplicons were assembled into plasmid PLAU36 linearised using HindIII and EcoRI via homologous recombination in *S. cerevisiae*.

A series of constructs were cloned containing various amounts of the additional sequence upstream of GFP which would correspond to the DNA present in the circular form of the element. This was achieved by first producing a construct containing 1,902 bp of additional sequence. The fragment containing the hygromycin-resistance cassette and the circular junction sequence of *Hephaestus* from the basic *starmid* construct was amplified with AD90+AD93 and the GFP coding region and terminator of plasmid PLAU17 was amplified with AD94+AD95 ^43^. These fragments were cloned into plasmid PLAU36 linearised using HindIII and EcoRI via homologous recombination in *S. cerevisiae* ^15^. From this base construct a series of constructs were produced by amplifying GFP along with variable amounts of upstream DNA using primers AD158 to AD166 in combination with primer AD67 and combining these products with the hygromycin resistance cassette amplified with primer pair AD90+AD67. These amplicons were assembled into plasmid PLAU36 linearised using HindIII and EcoRI via homologous recombination in *S. cerevisiae*. The resulting constructs contain 99 bp (AD158), 199 bp (AD159), 299 bp (AD160), 399 bp (AD161), 499 bp (AD162), 699 bp (AD163), 899 bp (AD164), 1,099 bp (AD165) and 1,489 bp (AD166) of “circular promoter” DNA upstream from the TTAC microhomologies which flank *Hephaestus*.

These constructs were transformed into the *Hephaestus-*negative *Paecilomyces* strain CBS 101075. This was accomplished via *Agrobacterium*-mediated transformation carried out exactly as described previously ^29^. Transformants were grown for 6 weeks at ambient temperature because at this point expression from the linear form of the promoter was low.

### Flow cytometry

WT and GFP-expressing spores were acquired on a 5-laser LSRFortessa cytometer equipped with Blue 488 nm and Yellow-green (YG) 561 nm lasers. Briefly, WT spores were used to set particle scatter and baseline fluorescence parameters for GFP (Blue 530/30 nm) and a co-visualisation detector (YG 586/15 nm). Tight, hierarchical gating selected single, consistently sized spores and at least 20,000 gated events were recorded. GFP median fluorescence intensity (MFI) values were batch processed using FCS Express 7 (De Novo software, Pasadena, CA). Three independent transformants were assayed for each construct. Median fluorescence intensity for each transformant was used for statistical analysis. Statistical significance between the short- and long-promoter groups was assessed using a two-sided unpaired t-test.

### Curing of strains carrying *starmids* using FdU selection

The loss of the thymidine kinase (TK) gene was selected for by embedding ∼10^7^ conidia in potato dextrose agar supplemented with 0.1 µM fluorodeoxyuridine (FdU). Spontaneously resistant colonies were passaged onto fresh media and replated onto hygromycin to identify a sensitive isolate in which the *starmid* had been lost.

### Testing for linked movement of *Hephaestus*

A mini-*Hephaestus* transformant of CBS 101075 previously generated ^15^ was crossed to *Paecilomyces* strain CBS 144490 by coculturing strains on PDA for approximately 4 weeks at 30°C. Material containing a mixture of sexual and asexual spores was resuspended in 200 µl of water using a pipette tip and incubated at 80°C for 10 minutes which is sufficient to kill the asexual but not the sexual fungal spores ^47^. Progeny were then plated onto PDA and colonies screened by PCR to identify one containing both the mini and full-length *Hephaestus*. Approximately 10^8^ conidia were plated onto 9 cm potato dextrose hygromycin plates and spontaneously hygromycin-resistant colonies selected (indicating transposition of the *mini-Hephaestus* element). Movement of the full-length *Hephaestus* in these strains was assayed by PCR using a rapid DNA extraction protocol ^48^ and Taq polymerase isolated from the pTaq plasmid ^49^. Primers used were AD304, AD305 and AD306 which gave a size shift in the amplified product from approximately 250 bp to 400 bp if *Hephaestus* transposes.

## Data Availability

Newly generated sequencing data have been deposited in NCBI BioProject PRJNA1532785.

## Acknowledgements

We are grateful to Alexander Idnurm for reading an earlier draft of this manuscript. ASU is supported by the Australian Research Council through a Discovery Early Career Researcher Award.

## Supplementary

**Figure S1.**
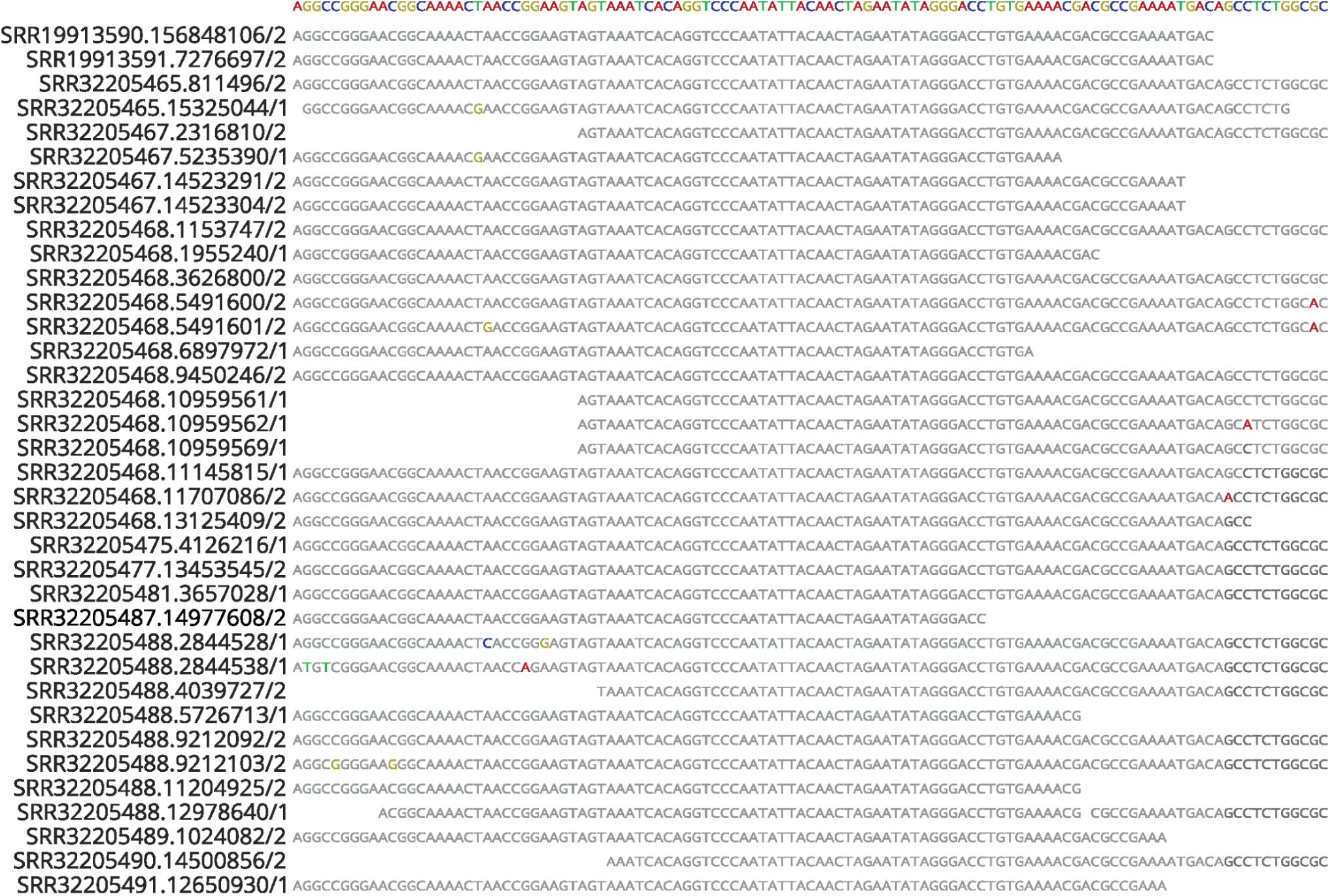
Illumina sequencing reads spanning the junction of the circular intermediate

**Table S1.** SRA accessions examined for reads indicative of the circular intermediate.

| SRA accession | Description | Gbp | Supporting reads |
| --- | --- | --- | --- |
| SRR8113285 | Genome sequencing of a T-DNA mutant of *Paecilomyces variotii* CBS 144490 | 3 |  |
| SRR19913590 | *Paecilomyces variotii* CBS 101075 transformed with a mini-*Hephaestus* construct. | 84.7 | 1 |
| SRR19913591 | *Paecilomyces variotii* CBS 101075 in which *Hephaestus* was flanked within a hygromycin resistance cassette. | 6.3 | 1 |
| SRR10993199 | *Paecilomyces variotii* FRR 3823 | 6.7 |  |
| SRR10993201 | *Paecilomyces variotii* FRR 1658 | 5.9 |  |
| SRR10993198 | *Paecilomyces paravariotii* FRR 5287 | 4.7 |  |
| SRR24236425 | *Paecilomyces variotii* DTO 021-C3 | 2.4 |  |
| SRR24236421 | *Paecilomyces variotii* DTO 027-B3 | 2.5 |  |
| SRR24236420 | *Paecilomyces variotii* DTO 027-B5 | 2.4 |  |
| SRR24236409 | *Paecilomyces variotii* DTO 169-C6 | 2.1 |  |
| SRR32217672 | *Paecilomyces variotii* CBS144490 | 5 |  |
| SRR32205493 | *Paecilomyces paravariotii* FRR 5287 | 5.5 |  |
| SRR32205490 | Recipient strain in *Hephaestus* transfer experiment | 4.9 | 1 |
| SRR32205487 | Recipient strain in *Hephaestus* transfer experiment | 4.5 | 1 |
| SRR32205484 | Recipient strain in *Hephaestus* transfer experiment | 5.2 |  |
| SRR32205481 | Recipient strain in *Hephaestus* transfer experiment | 3.5 | 1 |
| SRR32205478 | Recipient strain in *Hephaestus* transfer experiment | 5 |  |
| SRR32205489 | Recipient strain in *Hephaestus* transfer experiment | 4.6 | 1 |
| SRR32205486 | Recipient strain in *Hephaestus* transfer experiment | 4.5 |  |
| SRR32205483 | Recipient strain in *Hephaestus* transfer experiment | 4.8 |  |
| SRR32205480 | Recipient strain in *Hephaestus* transfer experiment | 5.2 |  |
| SRR32205475 | Recipient strain in *Hephaestus* transfer experiment | 5.5 | 1 |
| SRR32205477 | Recipient strain in *Hephaestus* transfer experiment | 5.1 | 1 |
| SRR32205474 | Recipient strain in *Hephaestus* transfer experiment | 5.3 |  |
| SRR32205472 | Recipient strain in *Hephaestus* transfer experiment | 5.3 |  |
| SRR32205469 | Recipient strain in *Hephaestus* transfer experiment | 3.8 |  |
| SRR32205471 | Recipient strain in *Hephaestus* transfer experiment | 5.4 |  |
| SRR32205468 | Recipient strain in *Hephaestus* transfer experiment | 4.2 | 13 |
| SRR32205466 | Recipient strain in *Hephaestus* transfer experiment | 5 |  |
| SRR32205465 | Recipient strain in *Hephaestus* transfer experiment | 4.8 | 1 |
| SRR32205491 | Recipient strain in *Hephaestus* transfer experiment | 4.9 | 1 |
| SRR32205488 | Recipient strain in *Hephaestus* transfer experiment | 5.1 | 8 |
| SRR32205485 | Recipient strain in *Hephaestus* transfer experiment | 5 |  |
| SRR32205482 | Recipient strain in *Hephaestus* transfer experiment | 4.7 |  |
| SRR32205479 | Recipient strain in *Hephaestus* transfer experiment | 4 |  |
| SRR32205476 | Recipient strain in *Hephaestus* transfer experiment | 4.3 |  |
| SRR32205473 | Recipient strain in *Hephaestus* transfer experiment | 5.2 |  |
| SRR32205470 | Recipient strain in *Hephaestus* transfer experiment | 3.2 |  |
| SRR32205467 | Recipient strain in *Hephaestus* transfer experiment | 4.7 | 4 |

**Table S2.**
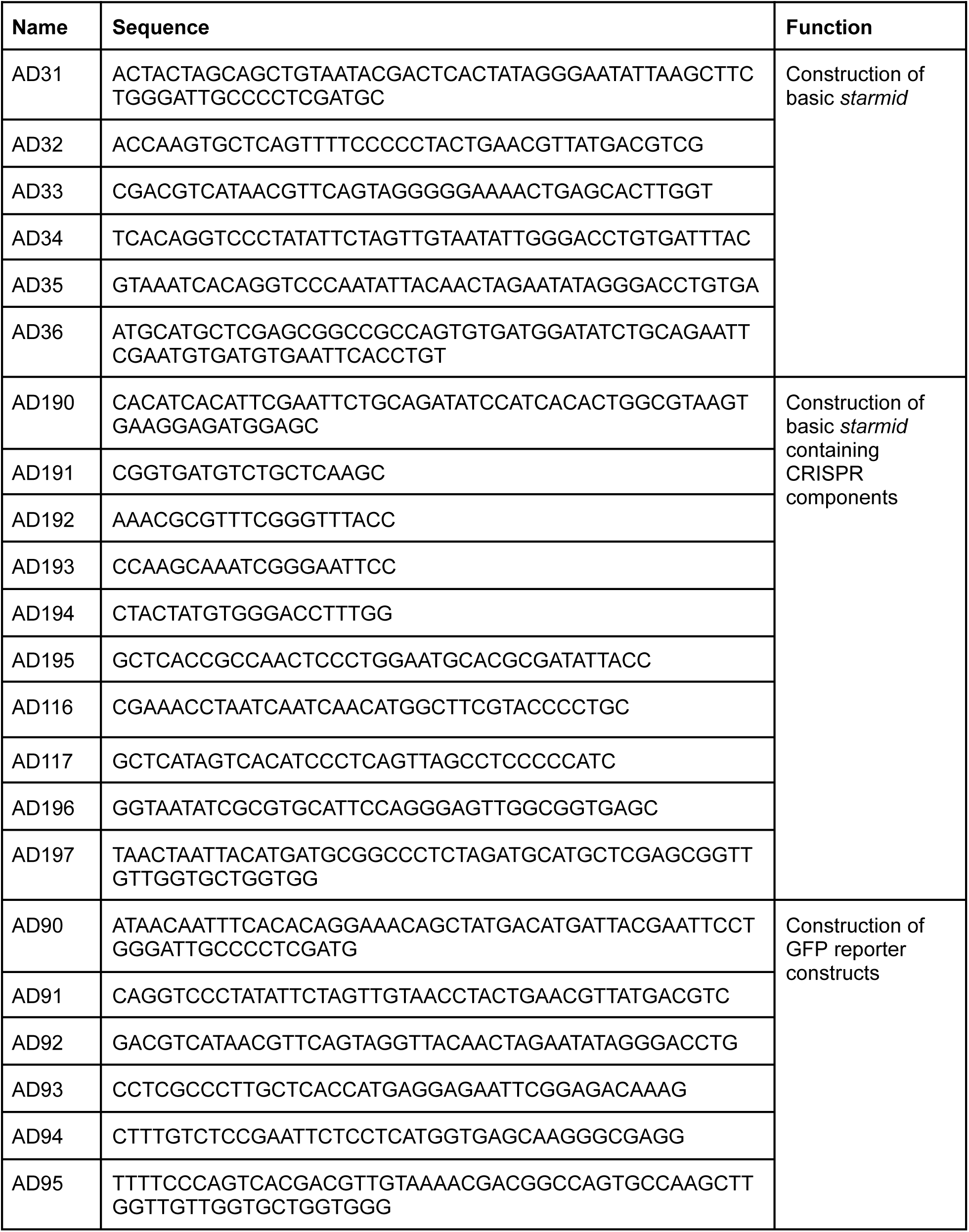
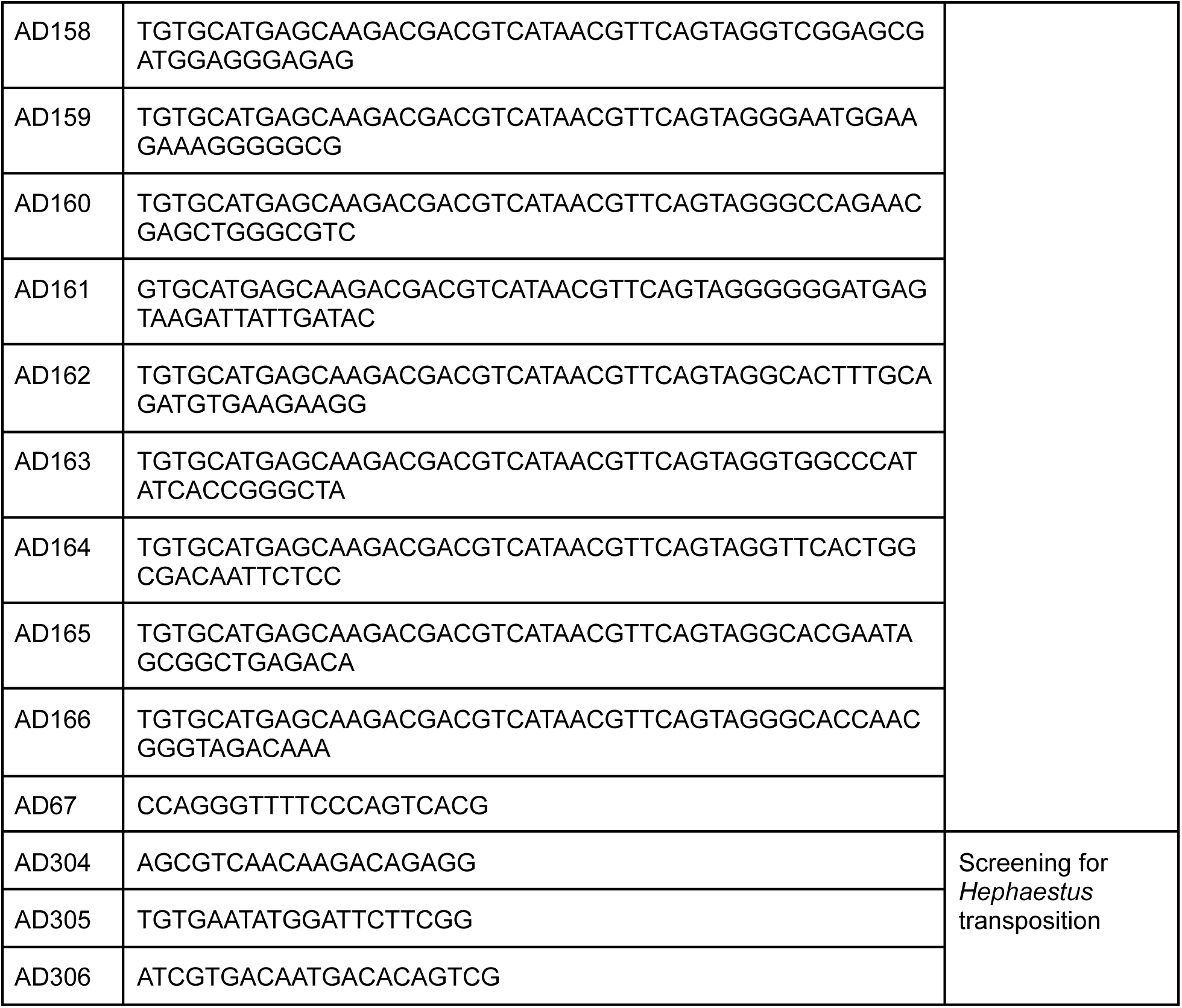
Oligonucleotides used in this study.

## References

1. Frost, L.S., Leplae, R., Summers, A.O., and Toussaint, A. (2005). Mobile genetic elements: the agents of open source evolution. Nat. Rev. Microbiol. 3, 722–732.

2. Ventola, C.L. (2015). The antibiotic resistance crisis: part 1: causes and threats. P T 40, 277–283.

3. Lander, E.S., Linton, L.M., Birren, B., Nusbaum, C., Zody, M.C., Baldwin, J., Devon, K., Dewar, K., Doyle, M., FitzHugh, W., et al. (2001). Initial sequencing and analysis of the human genome. Nature 409, 860–921.

4. De Cock, K.M., Jaffe, H.W., and Curran, J.W. (2021). Reflections on 40 years of AIDS. Emerg. Infect. Dis. 27, 1553–1560.

5. Cohen, S.N., Chang, A.C., Boyer, H.W., and Helling, R.B. (1973). Construction of biologically functional bacterial plasmids in vitro. Proc. Natl. Acad. Sci. U. S. A. 70, 3240–3244.

6. Jackson, D.A., Symons, R.H., and Berg, P. (1972). Biochemical method for inserting new genetic information into DNA of Simian Virus 40: circular SV40 DNA molecules containing lambda phage genes and the galactose operon of *Escherichia coli*. Proc. Natl. Acad. Sci. U. S. A. 69, 2904–2909.

7. Baltimore, D. (1970). RNA-dependent DNA polymerase in virions of RNA tumour viruses. Nature 226, 1209–1211.

8. Temin, H.M., and Mizutani, S. (1970). RNA-dependent DNA polymerase in virions of Rous sarcoma virus. Nature 226, 1211–1213.

9. Weiss, B., and Richardson, C.C. (1967). Enzymatic breakage and joining of deoxyribonucleic acid, I. Repair of single-strand breaks in DNA by an enzyme system from *Escherichia coli* infected with T4 bacteriophage. Proc. Natl. Acad. Sci. U. S. A. 57, 1021–1028.

10. Sternberg, N., and Hamilton, D. (1981). Bacteriophage P1 site-specific recombination. I. Recombination between *loxP* sites. J. Mol. Biol. 150, 467–486.

11. Smith, H.O., and Wilcox, K.W. (1970). A restriction enzyme from *Hemophilus influenzae*. I. Purification and general properties. J. Mol. Biol. 51, 379–391.

12. Jinek, M., Chylinski, K., Fonfara, I., Hauer, M., Doudna, J.A., and Charpentier, E. (2012). A programmable dual-RNA-guided DNA endonuclease in adaptive bacterial immunity. Science 337, 816–821.

13. McClintock, B. (1950). The origin and behavior of mutable loci in maize. Proc. Natl. Acad. Sci. U. S. A. 36, 344–355.

14. Urquhart, A.S., Chong, N.F., Yang, Y., and Idnurm, A. (2022). A large transposable element mediates metal resistance in the fungus *Paecilomyces variotii*. Curr. Biol. 32, 937–950.e5.

15. Urquhart, A.S., Vogan, A.A., Gardiner, D.M., and Idnurm, A. (2023). *Starships* are active eukaryotic transposable elements mobilized by a new family of tyrosine recombinases. Proc. Natl. Acad. Sci. U. S. A. 120, e2214521120.

16. Gluck-Thaler, E., Ralston, T., Konkel, Z., Ocampos, C.G., Ganeshan, V.D., Dorrance, A.E., Niblack, T.L., Wood, C.W., Slot, J.C., Lopez-Nicora, H.D., et al. (2022). Giant *Starship* elements mobilize accessory genes in fungal genomes. Mol. Biol. Evol. 39, msac109.

17. Urquhart, A., O’Donnell, S., Gluck-Thaler, E., and Vogan, A. (2025). A natural mechanism of eukaryotic horizontal gene transfer. bioRxiv, 10.1101/2025.02.28.640899.

18. Bucknell, A., Wilson, H.M., do Santos, K.C.G., Simpfendorfer, S., Milgate, A., Germain, H., Solomon, P.S., Bentham, A., and McDonald, M.C. (2025). *Sanctuary*: A *Starship* transposon facilitating the movement of the virulence factor ToxA in fungal wheat pathogens. MBio 16, e01371–25.

19. Sato, Y., Bex, R., van den Berg, G.C.M., Santhanam, P., Höfte, M., Seidl, M.F., and Thomma, B.P.H.J. (2025). *Starship* giant transposons dominate plastic genomic regions in a fungal plant pathogen and drive virulence evolution. Nat. Commun. 16, 6806.

20. Gluck-Thaler, E., Forsythe, A., Puerner, C., Gutierrez-Perez, C., Stajich, J.E., Croll, D., Cramer, R.A., and Vogan, A.A. (2025). Giant transposons promote strain heterogeneity in a major fungal pathogen. MBio 16, e0109225.

21. Urquhart, A.S., Forsythe, A., and Vogan, A.A. (2025). Are fungal disease outbreaks instigated by *Starship* transposons? Mol. Plant Pathol. 26, e70124.

22. Urquhart, A.S., Gluck-Thaler, E., and Vogan, A.A. (2024). Gene acquisition by giant transposons primes eukaryotes for rapid evolution via horizontal gene transfer. Sci. Adv. 10, eadp8738.

23. O’Donnell, S., Rezende, G., Vernadet, J.-P., Snirc, A., and Ropars, J. (2025). Harboring *Starships*: The accumulation of large horizontal gene transfers in domesticated and pathogenic fungi. Genome Biol. Evol. 17, evaf125.

24. Goodwin, T.J.D., Butler, M.I., and Poulter, R.T.M. (2003). Cryptons: a group of tyrosine-recombinase-encoding DNA transposons from pathogenic fungi. Microbiology 149, 3099–3109.

25. Kojima, K.K., and Jurka, J. (2011). Crypton transposons: identification of new diverse families and ancient domestication events. Mob. DNA 2, 12.

26. Nagy, A. (2000). Cre recombinase: the universal reagent for genome tailoring. Genesis 26, 99–109.

27. Wozniak, R.A.F., and Waldor, M.K. (2010). Integrative and conjugative elements: mosaic mobile genetic elements enabling dynamic lateral gene flow. Nat. Rev. Microbiol. 8, 552–563.

28. Bucknell, A.H., and McDonald, M.C. (2023). That’s no moon, it’s a *Starship*: Giant transposons driving fungal horizontal gene transfer. Mole. Microbiol. 120, 555–563.

29. Urquhart, A.S., Mondo, S.J., Mäkelä, M.R., Hane, J.K., Wiebenga, A., He, G., Mihaltcheva, S., Pangilinan, J., Lipzen, A., Barry, K., et al. (2018). Genomic and genetic insights into a cosmopolitan fungus, *Paecilomyces variotii* (Eurotiales). Front. Microbiol. 9, 3058.

30. van den Brule, T. (2022). Heterogeneity in stress resistance of Paecilomyces variotii and related food spoilage fungi. 10.33540/1362.

31. Urquhart, A., Vogan, A.A., and Gluck-Thaler, E. (2024). Starships: a new frontier for fungal biology. Trends Genet. 40, 1060–1073.

32. Urquhart, A.S., Hu, J., Chooi, Y.-H., and Idnurm, A. (2019). The fungal gene cluster for biosynthesis of the antibacterial agent viriditoxin. Fungal Biol. Biotechnol. 6, 2.

33. Gardiner, D.M., and Howlett, B.J. (2004). Negative selection using thymidine kinase increases the efficiency of recovery of transformants with targeted genes in the filamentous fungus *Leptosphaeria maculans*. Curr. Genet. 45, 249–255.

34. Hamari, Z., Amillis, S., Drevet, C., Apostolaki, A., Vágvölgyi, C., Diallinas, G., and Scazzocchio, C. (2009). Convergent evolution and orphan genes in the Fur4p-like family and characterization of a general nucleoside transporter in *Aspergillus nidulans*. Mol. Microbiol. 73, 43–57.

35. Sentchilo, V., Zehnder, A.J.B., and van der Meer, J.R. (2003). Characterization of two alternative promoters for integrase expression in the *clc* genomic island of *Pseudomonas* sp. strain B13. Mol. Microbiol. 49, 93–104.

36. Roux, I., and Chooi, Y.-H. (2022). Cre/*lox*-mediated chromosomal integration of biosynthetic gene clusters for heterologous expression in *Aspergillus nidulans*. ACS Synth. Biol. 11, 1186–1195.

37. Gluck-Thaler, E., and Vogan, A.A. (2024). Systematic identification of cargo-mobilizing genetic elements reveals new dimensions of eukaryotic diversity. Nucleic Acids Res. 52, 5496–5513.

38. Gems, D., Johnstone, I.L., and Clutterbuck, A.J. (1991). An autonomously replicating plasmid transforms *Aspergillus nidulans* at high frequency. Gene 98, 61–67.

39. Seekles, S.J., Teunisse, P.P.P., Punt, M., van den Brule, T., Dijksterhuis, J., Houbraken, J., Wösten, H.A.B., and Ram, A.F.J. (2021). Preservation stress resistance of melanin deficient conidia from *Paecilomyces variotii* and *Penicillium roqueforti* mutants generated via CRISPR/Cas9 genome editing. Fungal Biol. and Biotechnol. 8, 4.

40. Han, H.-G., Nandre, R., Eom, H., Choi, Y.-J., and Ro, H.-S. (2025). Development of a CRISPR/Cas9 RNP-mediated genetic engineering system in *Paecilomyces* variotii. J. Microbiol. 63, e2502011.

41. van Rhijn, N., Furukawa, T., Zhao, C., McCann, B.L., Bignell, E., and Bromley, M.J. (2020). Development of a marker-free mutagenesis system using CRISPR-Cas9 in the pathogenic mould *Aspergillus fumigatus*. Fungal Genet. Biol. 145, 103479.

42. Langmead, B., and Salzberg, S.L. (2012). Fast gapped-read alignment with Bowtie 2. Nat. Methods 9, 357–359.

43. Idnurm, A., Urquhart, A.S., Vummadi, D.R., Chang, S., Van de Wouw, A.P., and López-Ruiz, F.J. (2017). Spontaneous and CRISPR/Cas9-induced mutation of the osmosensor histidine kinase of the canola pathogen *Leptosphaeria maculans*. Fungal Biol. Biotechnol. 4, 12.

44. Nødvig, C.S., Nielsen, J.B., Kogle, M.E., and Mortensen, U.H. (2015). A CRISPR-Cas9 system for genetic engineering of filamentous fungi. PLoS One 10, e0133085.

45. Arentshorst, M., Ram, A.F.J., and Meyer, V. (2012). Using non-homologous end-joining-deficient strains for functional gene analyses in filamentous fungi. In Plant Fungal Pathogens: Methods and Protocols Methods in molecular biology (Clifton, N.J.). (Humana Press), pp. 133–150.

46. Pitkin, J.W., Panaccione, D.G., and Walton, J.D. (1996). A putative cyclic peptide efflux pump encoded by the TOXA gene of the plant-pathogenic fungus Cochliobolus carbonum. Microbiology 142 ( *Pt 6*), 1557–1565.

47. Houbraken, J., Varga, J., Rico-Munoz, E., Johnson, S., and Samson, R.A. (2008). Sexual reproduction as the cause of heat resistance in the food spoilage fungus *Byssochlamys spectabilis* (anamorph *Paecilomyces variotii)*. Appl. Environ. Microbiol. 74, 1613–1619.

48. Liu, K.-H., Yeh, Y.-L., and Shen, W.-C. (2011). Fast preparation of fungal DNA for PCR screening. J. Microbiol. Methods 85, 170–172.

49. Desai, U.J., and Pfaffle, P.K. (1995). Single-step purification of a thermostable DNA polymerase expressed in *Escherichia coli*. Biotechniques 19, 780–782, 784.

